# Recovery and Community Succession of the *Zostera marina* Rhizobiome After Transplantation

**DOI:** 10.1101/2020.04.20.052357

**Authors:** Lu Wang, Mary K. English, Fiona Tomas, Ryan S. Mueller

**Affiliations:** Department of Microbiology, Oregon State University, Corvallis, OR, USA; Instituto Mediterráneo de Estudios Avanzados (CSIC-UIB), Balearic Islands, Spain; Department of Fisheries and Wildlife, Oregon State University, Corvallis, OR, USA

**Author notes:** Address correspondence to Ryan S. Mueller,.

**Keywords:** seagrass, eelgrass, restoration, rhizosphere, rhizoplane, sulfur-oxidizing bacteria, cable bacteria, microbial diversity, succession

## Abstract

Seagrasses can form mutualisms with their microbiomes that facilitate the exchange of energy sources, nutrients, and hormones, and ultimately impact plant stress resistance. Little is known about community succession within the belowground seagrass microbiome after disturbance and its potential role in the plant’s recovery after transplantation. We transplanted *Zostera marina* shoots with and without an intact rhizosphere and cultivated plants for four weeks while characterizing microbiome recovery and effects on plant traits. Rhizosphere and root microbiomes were compositionally distinct, likely representing discrete microbial niches. Furthermore, microbiomes of washed transplants were initially different from those of sod transplants, and recovered to resemble an undisturbed state within fourteen days. Conspicuously, changes in microbial communities of washed transplants corresponded with changes in rhizosphere sediment mass and root biomass, highlighting the strength and responsive nature of the relationship between plants, their microbiome, and the environment. Potential mutualistic microbes that were enriched over time include those that function in the cycling and turnover of sulfur, nitrogen, and plant-derived carbon in the rhizosphere environment. These findings highlight the importance and resiliency of the seagrass microbiome after disturbance. Consideration of the microbiome will have meaningful implications on habitat restoration practices.

**Importance:** Seagrasses are important coastal species that are declining globally, and transplantation can be used to combat these declines. However, the bacterial communities associated with seagrass rhizospheres and roots (the microbiome) are often disturbed or removed completely prior to transplantation. The seagrass microbiome benefits seagrasses through metabolite, nutrient, and phytohormone exchange, and contributes to the ecosystem services of seagrass meadows by cycling sulfur, nitrogen, and carbon. This experiment aimed to characterize the importance and resilience of the seagrass belowground microbiome by transplanting *Zostera marina* with and without intact rhizospheres and tracking microbiome and plant morphological recovery over four weeks. We found the seagrass microbiome to be resilient to transplantation disturbance, recovering after fourteen days. Additionally, microbiome recovery was linked with seagrass morphology, coinciding with increases in rhizosphere sediment mass and root biomass. Results of this study can be used to include microbiome responses in informing future restoration work.

## Introduction

The rhizobiome has long been recognized to have important impacts on plant growth and health (1). The microbes of the rhizobiome, which directly interact with and are influenced by the roots (2), can benefit their plant hosts through recycling and producing bioavailable nutrients (3–5), increasing disease resistance through competition with or inhibition of pathogens (6), and influencing plant growth and stress tolerance through production of phytohormones (7, 8). Community composition within the rhizobiome is shaped by plant metabolism and physiology, which controls rhizodeposition, exudation of organic carbon and nitrogen, and release of defense compounds (7, 9, 10). The quantity and composition of exudates can impact microbial activity in the rhizosphere and vary as a result of many factors (11–14). While plant-rhizobiome interactions are relatively well-defined for terrestrial plants, analogous interactions between aquatic plants and their microbiomes have only recently started to become known (15, 16).

Seagrasses are marine vascular plants that form key ecosystems on coastal areas worldwide, where they provide numerous ecosystem services (17). Recent evidence suggests that members of the seagrass microbiome may modulate host growth and response to environmental stresses (15, 18, 19). In addition to fixing nitrogen and producing phytohormones (20, 21), the seagrass microbiome is proposed to mitigate the toxic effects of hydrogen sulfide in sediments, which have been linked to declines in seagrass health and localized die-back events (22–24). The seagrass rhizobiome is thought to be primarily influenced by exudation of carbon compounds, which can provide up to 60% of the carbon assimilated by these microbes (25, 26), and by radial oxygen loss from roots, which may promote colonization of the rhizosphere by distinct bacteria (24, 27).

The effect of rhizosphere disturbance on the composition of seagrass microbiomes and plant health has rarely been explored (28). Yet, it may be important both for plant recovery after a disturbance and in the context of restoration outcomes, which are highly variable and dependent on methodology (29–32). Sod transplants, which transfer shoots with intact rhizospheres, have historically been one of the more successful methods, potentially because the intact rhizosphere sediment acts as a natural anchor and retains functional relationships between the plant and its rhizobiome (31). Conversely, bare root transplants are generally less successful and could experience a decrease or lag in plant performance as the rhizobiome redevelops after transplantation. Importantly, microbial community succession after disturbance can strongly affect host health in several microbiome-host systems (e.g., algae, corals, and humans), whereby dysbiosis disrupts host functioning and increases susceptibility to disease (33–35). Thus, it is important to understand the recovery of seagrass microbiomes after disturbance, as this may impact seagrass health and resistance to environmental stresses.

In this study, we characterized the recovery of seagrass rhizobiomes post-disturbance by transplanting *Zostera marina* with and without an intact rhizosphere and sampling for plant and microbiome characteristics over the course of 28 days. We expected to see the rhizobiome of seagrass transplanted without an intact rhizosphere recover over time to resemble that of the control plants, with a corresponding delay in the response of plant growth traits.

## Results

### *Changes in* Z. marina *Traits After Transplantation*

We quantified several traits to assess plant growth and measure the rhizosphere sediment mass recovery on roots after transplantation (Table S2). Plant traits varied significantly due to an interaction between days post transplantation (DPT) and treatment (PERMANOVA: DPT x Treatment *F*_*1,61*_ = 2.85, *p* = .036, *R*^*2*^ = .03; Table S3). While plant traits did not differ amongst treatments at the beginning of the experiment (PERMANOVA: Day 0 Treatment *F*_*1,8*_ = 1.12, *p* = .304, *R*^*2*^ = .12; Figures 1B and S1), they exhibited overall differences within seven days after transplantation, and plant traits of the wash treatment began to more strongly resemble those of the sod treatment after one week (Figure 1A). For sod transplants, the most variation in traits occurred within the first seven days of the experiment, after which these measures stabilized and remained relatively constant (Figures 1B and S1). Conversely, changes in the traits of washed plants occurred more slowly, stabilizing only after fourteen days. By the end of the experiment no between-treatment variation in traits was evident (PERMANOVA: Day 28 Treatment *F*_*1,13*_ = 1.00, *p* = .422, *R*^*2*^ = .07; Figures 1B and S1).

**Figure 1:**
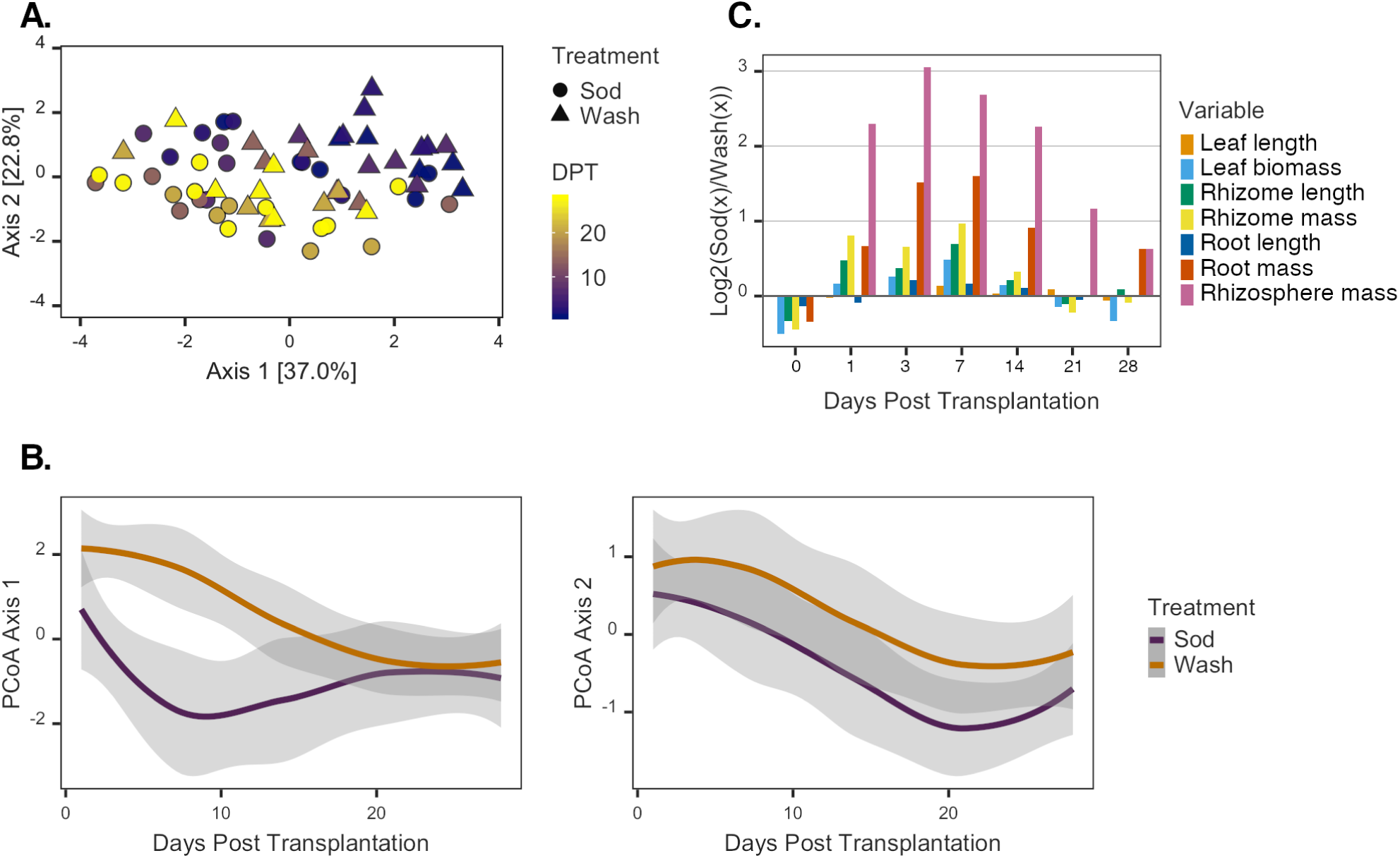
Variance in *Z. marina* Traits Over Time. (A) PCoA of *Z. marina* plants based on a Euclidean distance matrix relating plant traits. Color gradient represents the day of plant collection (DPT), and symbols represent the treatment assignment of each plant. (B) Loess smoothed estimates of the first two principal coordinate summary variables over time (cumulative variance = 59.8%). Shaded areas represent 95% confidence intervals of estimates, and colors designate each respective treatment (purple = sod transplants, gold = washed transplants). (C) Relative differences in log transformed values of *Z. marina* morphometric data at sampling points over time. Positive values indicate higher values in sod transplants than washed, and negative values indicate the opposite.

Most traits exhibited significant increases over the course of the experiment in both the wash and the sod treatment groups, indicating overall growth of *Z. marina* shoots after transplantation regardless of rhizosphere presence (Figure S1 & Tables S4-S8). For instance, upon experiment completion, total biomass of transplants had increased 1.5-fold on average, and lengths of leaves and rhizomes had increased 1.5 and 1.8-fold, respectively (Figure S1). Whereas differences in traits due to treatment were minimal at the beginning and end of the experiment, they were most pronounced from days one to fourteen of the experiment when sod transplants consistently demonstrated greater increases compared to those of the washed transplants (Figure 1C). For example, root biomass and root length were not significantly affected by rhizosphere removal at the beginning of the experiment (Student’s t-test [root biomass]: Wash M = 0.016 ± 0.012 g, Sod M = 0.012 ± 0.006 g, *t*(8) = -0.57, *p =* .58; Student’s t-test [root length]: Wash M = 6.04 ± 1.91 cm, Sod M = 4.46 ± 0.92 cm, *t*(8) = -1.67, *p =* .13). Importantly, though, sod transplants increased 1.7-fold in root biomass on average, whereas wash transplants increased 1.1-fold by the end of the experiment (ANCOVA: Treatment *F*_*1,72*_ = 16.16, *p* = .0001; Figures 1C & S1C, Table S7). As expected from our treatment, rhizosphere sediment mass significantly varied with the interaction between the time covariate and the main treatment effect (ANCOVA: DPT x Treatment *F*_*1,71*_ = 18.78, *p* < .00005; Figure S1D & Table S8). The rhizosphere mass attached to roots of sod transplants did not change significantly during the experiment, whereas sediment accumulation on washed roots rapidly increased after seven days post transplantation and recovered to levels observed on sod transplants by the end of the experiment (Welch’s t-test: Wash M= 16.34 ± 10.03 g, Sod M = 25.25 ± 13.26 g, *t*(13) = 1.48, *p =* .16, Figure S1D).

### *Microbial Community Differences Between* Z. marina *Rhizosphere and Roots*

When considering all samples, microbial communities were most strongly clustered based on compartment (PERMANOVA: Compartment *F*_*1,112*_ = 26.33, *p* = .001, *R*^*2*^ = .16; Figure 2A and Table S9). Forty-two prokaryotic ASVs exhibited significantly different relative abundances in the rhizosphere versus roots (Table S10). Twenty-five were enriched in the rhizosphere, while the remaining 17 were in greater relative abundance on roots (Figure 2B). Significant ASVs were most commonly assigned to the Proteobacteria and Bacteroidetes phyla (n = 18 and 12, respectively), with 66% of the former taxon and 75% of the latter detected in higher relative abundance in rhizosphere over root communities. Conversely, ASVs of the Epsilonbacteraeota phylum were typically in higher relative abundances in root samples (five of seven ASVs). Due to the strong effect of compartment on microbial community structure, the remaining microbial diversity results are presented separately for rhizosphere and root samples.

**Figure 2:**
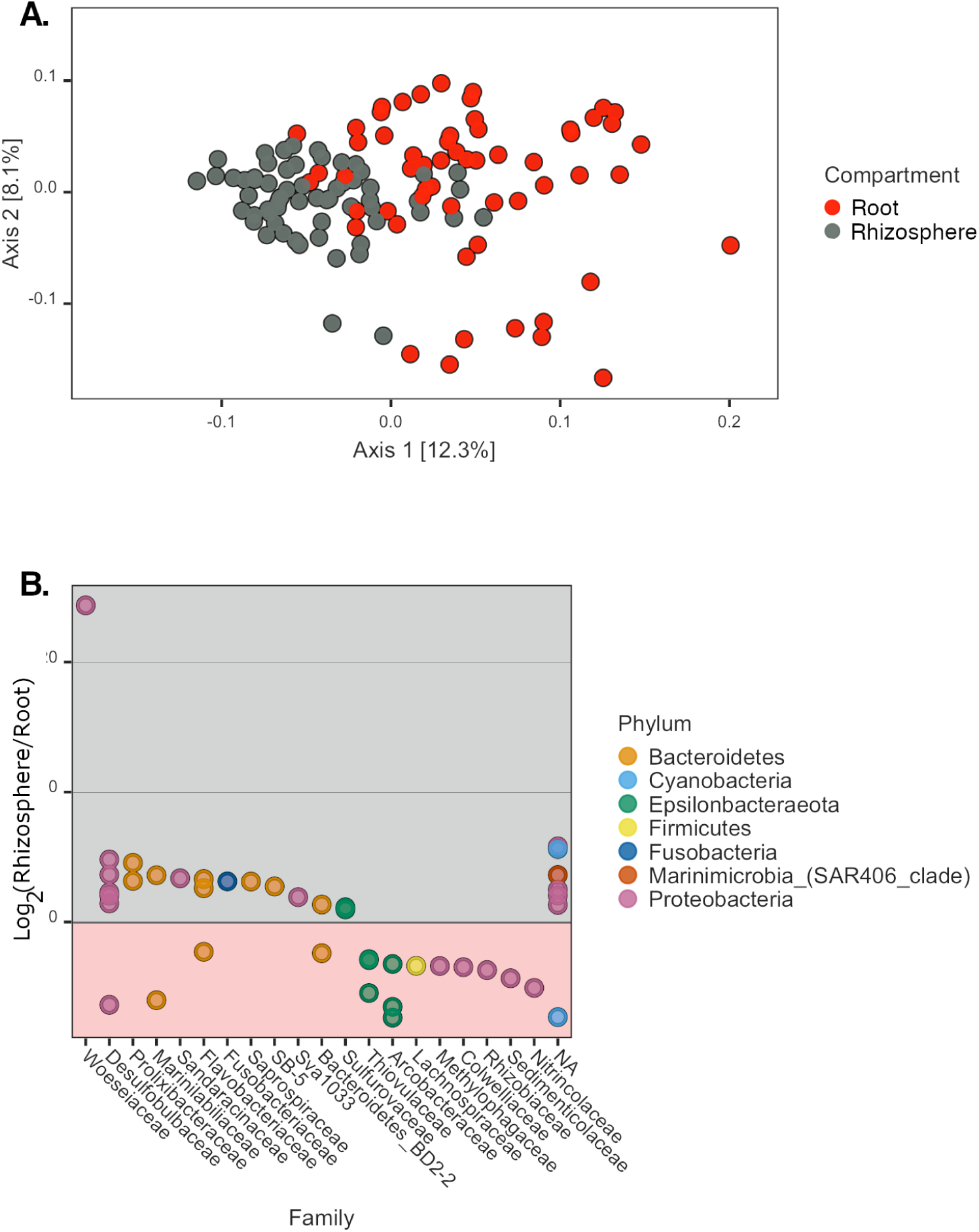
Microbial Community Differences Between *Z. marina* Compartments. (A) PCoA of *Z. marina* all sampled microbial communities based on a weighted UNIFRAC distance matrix. Colors indicate the compartment of each sample (Red = Root, Gray = Rhizosphere). (B) Taxa with significant relative abundance differences between compartments. Positive values indicate higher relative abundances of ASVs in rhizospheres than roots, and negative values indicate the opposite. ASVs assigned to the same phylum have the same color. ASVs are grouped by column by taxonomic family.

### Changes in Rhizosphere Microbiomes After Transplantation

Temporal changes in the structure of rhizosphere microbial communities mirror the patterns observed for plant trait data. That is, initial differences were observed between rhizosphere communities from plants of different treatment groups, but communities became more similar in structure by the end of the experiment (Figure 3A). The most variation was due to a shift of rhizosphere communities of washed transplants along the first principle coordinate to more strongly resemble sod samples after seven days. As observed for plant traits, a significant interactive effect of treatment and time on the rhizosphere community structure was detected (PERMANOVA: DPT x Treatment *F*_*1,58*_ = 2.53, *p* = .005, *R*^*2*^ = .03; Table S11).

**Figure 3:**
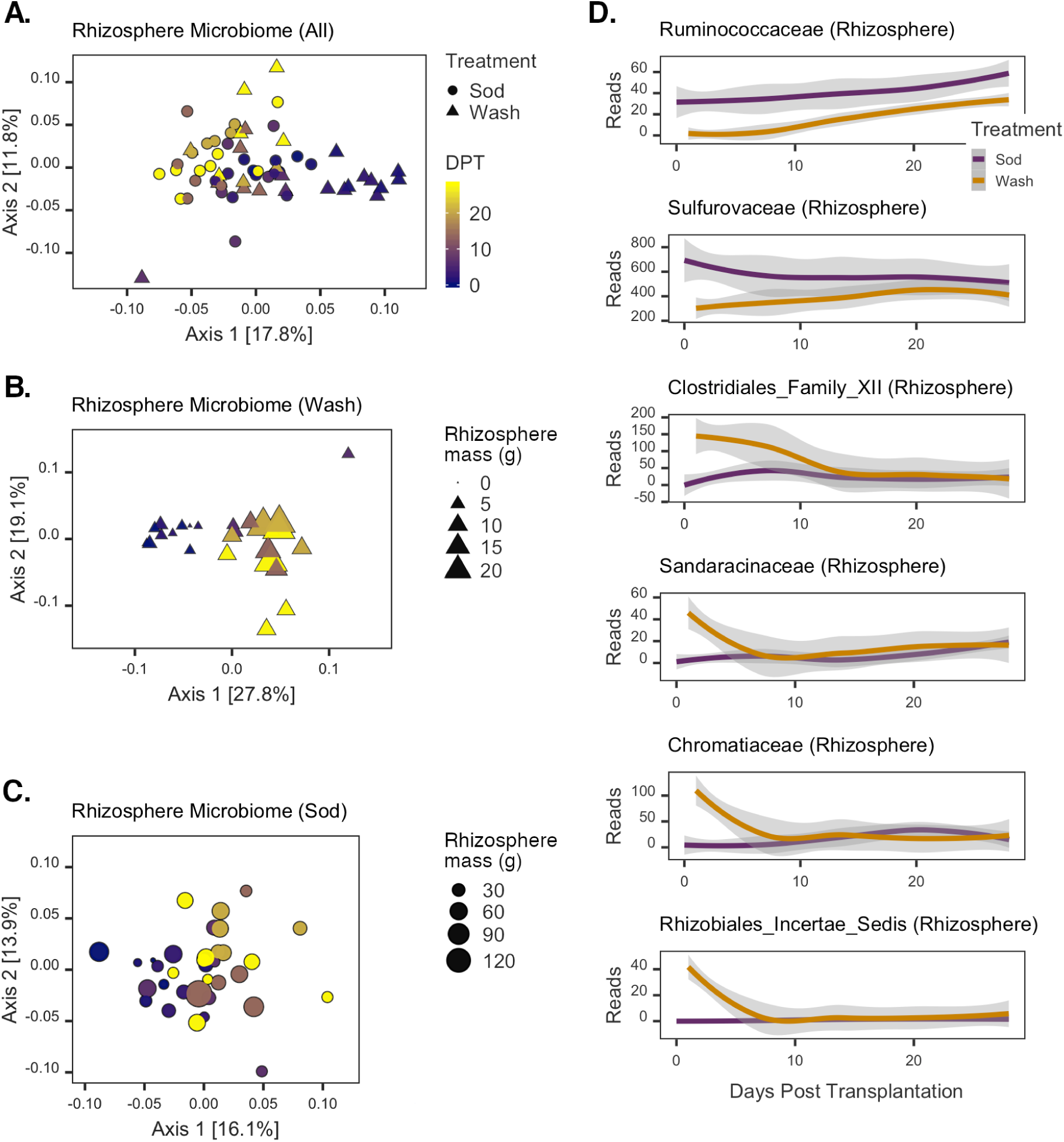
Changes in Rhizosphere Microbial Communities Post transplantation. PCoAs of (A) all rhizosphere, (B) washed rhizosphere, and (C) sod transplant rhizosphere communities. Color gradient represents the day of sample collection (DPT), and symbols represent the treatment assignment of each sample. Symbol size in (B) and (C) is scaled to the grams of rhizosphere sediment collected from each corresponding sampled plant. (D) Rhizosphere ASVs with significant Time x Treatment interaction effects. Lines are loess smoothed estimates of the sequence counts for each taxon; shaded areas represent 95% confidence intervals of estimates. Colors designate each respective treatment group (purple = sod transplants, gold = washed transplants).

To further investigate the treatment effect of rhizosphere disruption on the recovery of the rhizosphere communities, we analyzed the different treatment samples separately. Structural changes in the rhizosphere communities of the sod and wash treatment groups both demonstrated significant time effects, but a stronger temporal correlation was detected for the washed than sod transplant rhizosphere communities (PERMANOVA: DPT [Wash transplants] *F*_*1,25*_ = 6.47, *p* = 001, *R*^*2*^ = .21; DPT [Sod transplants] *F*_*1,33*_ = 3.57, *p* = .001, *R*^*2*^ = .10; Tables S12 & S13). A shift in community structure of washed transplants occurred at seven days and corresponded to the point of accelerating sediment accumulation on washed roots. Additionally, overall community changes were significantly correlated to rhizosphere sediment masses of all washed plants (Mantel test *p* = .004, Spearman’s *ρ* = .25; Figure 3B). For sod transplants, however, sediment mass was not correlated with rhizosphere community structure (Mantel test *p* = .43, Spearman’s *ρ* = .0001; Figure 3C), and was instead most strongly correlated with plant growth traits (Mantel test: Leaf Length + Rhizome Length + Leaf Biomass *p* = .001, Spearman’s *ρ* = .27).

We used regression analyses to identify microbial taxa that were specifically associated with *Z. marina* rhizosphere development during the experiment (GLMM: *adjusted p* ≤ .05; Table S14). Thirty-two taxonomic families had significantly different modeled intercepts between treatments, and 14 taxa exhibited significant differences in modeled slopes. Six taxa were found with significant differences in both slopes and intercepts (Figure 3D). Of these six, the Ruminococcaceae and Sulfurovaceae had negative intercepts and positive slope coefficients. For example, higher relative abundances of the Ruminococcaceae were detected in the sod samples on average, but the rate of increase of this taxon’s abundance was greater in washed samples over time. The Sulfurovaceae showed a similar temporal pattern of abundance in washed samples, but in sod transplants this taxon generally demonstrated a decrease over time. The remaining four taxa (Clostridiales Family XII, Sandaracinaceae, Chromatiaceae, and Rhizobiales [Incertae sedis]) all showed similar patterns (Figure 3D); in washed transplants they rapidly decreased to low levels within the first seven days of the experiment, whereas in sod transplants there was little to no detection of them throughout the experiment.

### Changes in Root Microbiomes After Transplantation

Recovery dynamics of root microbiomes were largely similar to those observed for rhizosphere communities (Figure 4A). A significant effect of the interaction between time and treatment on the structure of all root communities was detected (PERMANOVA: DPT x Treatment *F*_*1,58*_ = 2.01, *p* = .043, *R*^*2*^ = .03; Table S15). A relatively strong effect of time was evident for communities from wash transplants (PERMANOVA: *F*_*1,26*_ = 5.91, *p* = .001, *R*^*2*^ = .19; Figure 4B & Table S16), but not for sod transplants (PERMANOVA: *F*_*1,28*_ = 1.78, *p* = .096, *R*^*2*^ = .06; Figure 4C & Table S17). Changes in washed root microbiome community structure were not correlated with sediment mass accumulation (Mantel test: *p* = .085, Spearman’s *ρ* = .13), and were instead most strongly correlated with leaf length and rhizome mass (Mantel test: *p* = .001, Spearman’s *ρ* = .32). In contrast, the root microbiomes of sod transplants were relatively stable over time (PERMANOVA: *F*_*1,28*_ = 1.78, *p* = .096, *R*^*2*^ = .06; Figure 4C & Table S17), and not correlated with any single plant trait or combination thereof.

**Figure 4:**
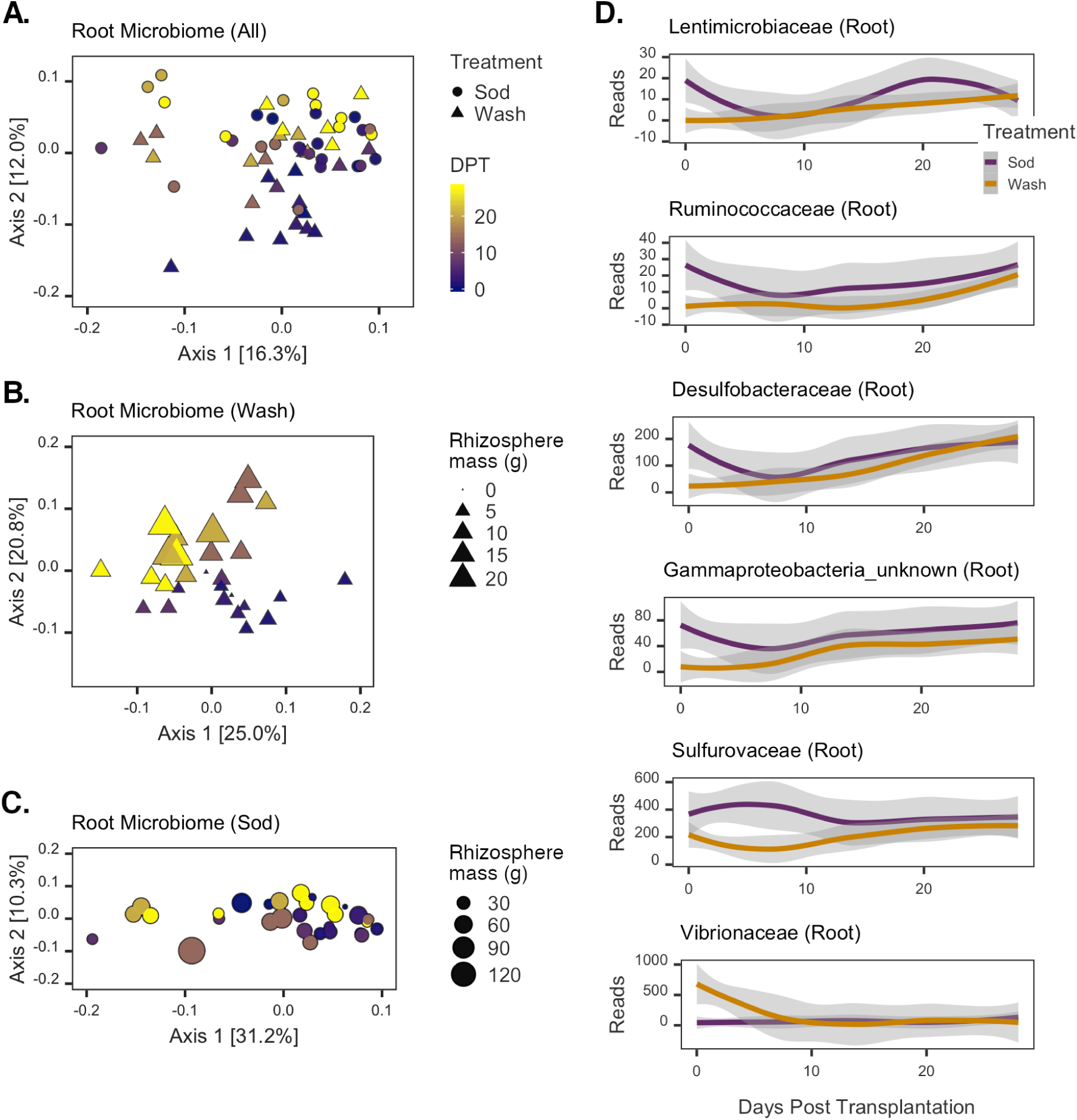
Changes in Root Microbial Communities Post transplantation. PCoAs of (A) all root, (B) washed root, and (C) sod transplant root communities. Color gradient represents the day of sample collection (DPT), and symbols represent the treatment assignment of each sample. Symbol size in (B) and (C) is scaled to the grams of rhizosphere sediment collected from each corresponding sampled plant. (D) Root ASVs with significant Time x Treatment interaction effects. Lines are loess smoothed estimates of the sequence counts for each taxon; shaded areas represent 95% confidence intervals of estimates. Colors designate each respective treatment group (purple = sod transplants, gold = washed transplants).

Regression analyses identified 25 taxa with significant differences in modeled intercepts between treatments, but no differences in modeled slopes (Table S18). Another four taxa were found to have no detectable differences in intercepts, but significant differences in slopes. Six taxa were found to have significant differences in both modeled intercepts and slopes (Figure 4D). The Lentimicrobiaceae, Ruminococcaceae, Desulfobacteraceae, and an unknown Gammaproteobacteria family were all modeled to have largely similar dynamics, with negative intercepts and positive slope coefficients. Abundances of these taxa on roots of sod transplants rapidly declined within seven days of transplantation, followed by a more gradual increase in abundance over the last two weeks of the experiment (Figure 4D). Conversely, these taxa were nearly undetectable initially on roots of washed transplants, but their abundances recovered by experiment completion. The Sulfurovaceae also exhibited gradual increases in relative abundance on washed roots, but in sod transplants this taxon’s abundance increased after seven days and subsequently decreased (Figure 4D). Vibrionaceae showed an altogether different pattern; high abundances were detected in initial wash samples, but by day seven these taxa were rarely detected. In sod transplants, this group was generally absent throughout the entire experiment.

## Discussion

Our results indicate that *Zostera marina* rhizobiome communities are distinct, linked to seagrass performance, and resilient to disturbance. Indeed, eelgrass belowground traits suffered negatively from undergoing rhizosphere removal, but they recovered after ca. 2 weeks. Concomitantly, their microbial communities resembled those of sod transplants by experiment end, indicating that *Z. marina* and its belowground microbiome are resilient to stresses associated with transplantation.

The observation of consistently distinct microbial communities between compartments of *Z. marina* is in line with studies describing the structure of seagrass microbiomes from field-collected samples, where large differences are observed between plant microbial communities and those in the surrounding environment (15, 18, 19, 36–39). In the study by Cúcio et al. (15) where the rhizosphere compartment was specifically analyzed, significant differences were found between communities of bulk and rhizosphere sediments. Our work further distinguishes the root-attached microbiome as different from the microbiota of the rhizosphere, and suggests that these two compartments are separate microbial niches shaped by prevailing redox and nutrient gradients formed across sub-millimeter ranges by plant metabolic processes (40). These results are supported by previous observations (19, Wang et al., submitted for publication) and a proposed model of microbiome assembly via selection of bulk sediment microbes (19).

Although the mechanisms controlling assembly of seagrass microbiomes are largely unknown, evidence from terrestrial plant studies suggest that they are based on metabolic interactions and nutrient exchange between plants and microbes. For instance, changes in abiotic factors and/or the presence of pathogens can induce or restrict exudation of nutritional and allelopathic compounds, contributing to the selection of a root microbiome (9, 41). Root exudation is known to be metabolically costly for plants, though, and can result in significant losses of carbon and nitrogen (7). However, these costs are likely offset by the beneficial functions of the belowground microbiome (e.g., disease suppression, nutrient acquisition, stress tolerance, and growth enhancement) (7, 8, 42).

Similar to these terrestrial plant examples, we propose that exudation is an important factor modulating belowground microbiomes of seagrasses. Seagrass exudation is known to change with environmental conditions (e.g. light restriction) (43), and can act as an important resource for sediment microbes (26, 44). Our concomitant observations of belowground biomass loss in washed *Z. marina* plants and large-scale changes in the microbiome structure within the first week after transplantation may be related to changes in root exudation, and would suggest a rapid and coordinated response by both the microbiome and plant to disturbance. While we cannot rule out the effects of potential root damage that may have occurred during seawater rinses to remove the rhizosphere prior to transplantation on microbiome community structure, our data suggest that no observable and significant root breakage and/or biomass loss occurred from initial washes.

When considering the timing of recovery between belowground compartments, it is notable that the change in microbial community structure of washed roots was detected three days after transplantation, whereas a similar change in the rhizosphere was detected on day seven (Figures 3B & 4B). This supports a possible role of root exudates in attracting or repelling microbial populations based on their proximity to the root surface. Interestingly, almost all of the root- and rhizosphere-associated taxa that significantly changed in abundance over time demonstrated an inflection point in their abundance trajectories between three and seven days post transplantation (Figures 3D & 4D). When considered with the changes observed for plant traits, these data suggest that the first week after transplantation is a critical transition period for the plant and its associated microbiome.

Rapid and resilient responses of microbiomes to disturbance have also been observed for microbiomes of terrestrial plants (45) and marine algae (34). In the latter study, community assembly on the surface of *Delisia pulchra* was found to be deterministic, recovering to a pre-disturbed state within 12 days. In this system, the production of anti-fouling chemicals (i.e., halogenated furanones) either by early-colonizing bacteria or by the algae, is an important factor controlling community succession. Seagrasses can also produce a diverse set of anti-fouling chemicals on their surfaces (46). Their precise role in modulating the epibiont community structure is currently unknown, but we believe that the collective results of ours and the aforementioned studies suggest that these compounds and nutritional exudates act to deterministically shape microbiome community structure.

Further, our results show that several taxa that may benefit the plant directly or enhance turnover of nutrients in sediments are enriched in seagrass-associated compartments after transplantation. ASVs assigned to the Bacteroidetes and the Sandaracinaceae taxa were always found to be significantly higher in relative abundance in rhizosphere over root samples. Both taxa are common inhabitants of marine environments and widely recognized to be important degraders of complex organic material, such as rhizodeposits, from plants (47–50). Notable taxa that were enriched on roots include ASVs with potential important roles in turnover of plant exudates. Namely, the Methylophagaceae are methanol consumers that can significantly impact plant growth and stress tolerance through the production of plant hormones (51, 52). Additionally, ASVs of the Lachnospiraceae and Colwelliaceae families that were enriched on roots may have potential roles in consumption of plant-derived polysaccharides and lignin (53, 54). In fact, the former group may have an additional symbiotic role with plants, as a novel species of Lachnospiraceae is proposed to be diazotrophic (55).

Other taxa found enriched in either the root or rhizosphere compartment appear to rely on respiratory metabolisms linked to sulfur and nitrogen cycles, a common feature of populations of the seagrass microbiome (20, 56). For example, lineages of the Desulfobulbaceae, which were more commonly enriched in the rhizosphere compartment, can act as strictly anaerobic sulfate reducers (e.g., *Desulforhopalus sp*.) (57), or as sulfide oxidizers whose filamentous cells respire by transferring electrons from reduced sulfur compounds across redox gradients to either oxygen or nitrate (e.g., the so-called ‘cable bacteria’ of *Ca. Electrothrix sp*.) (58, 59). When associated with *Z. marina*, Desulfobulbaceae cells may thrive in or around the oxic/anoxic transition zone within the rhizosphere where they transfer electrons to and from reduced sulfur compounds found within sediments. Such a preference is supported by recent work showing increased detection of these cells in low oxygen zones of seagrass roots (24) and the oxic-anoxic transition zone around roots of other aquatic plants (60). Several ASVs found enriched on roots were designated as known or putative sulfur-oxidizing bacteria (SOB), including Sedimenticolaceae (61), Thiovulaceae, and Arcobacteraceae (62), supporting the hypothesis that seagrasses facilitate the activities of SOB as a way to combat sulfide toxicity (63).

The Ruminococcaceae and the Sulfurovaceae stand out in our time-course analyses, as both exhibited similar abundance differences initially and over time in both root and rhizosphere samples. ASVs of these taxa, along with those of the Lentimicrobiaceae, Desulfobacteraceae, and Unknown Gammaproteobacteria, were noticeably absent on washed roots at the start of the experiment, but all recovered to the relatively high levels found on roots of sod transplants by the end of the experiment. Many of these taxa are known to drive sulfur cycling in marine sediments (62, 64, 65) and their functional roles may be important in long-term associations with plants. In contrast, the Vibrionaceae were the only taxa that rapidly decreased from high relative abundances on washed roots to undetectable levels after seven days. Given that many *Vibrio* species are pathogens that rapidly form biofilms on marine surfaces (66, 67) it is possible that these ASVs are detrimental to root health, and plants respond by changing chemical exudation as a way to discourage growth of these of bacteria while encouraging growth of beneficial microbes shortly after disturbance, when plants are vulnerable to transplantation stresses (i.e., transplantation shock).

Seagrass health after transplantation is often unpredictable (30) and restoration success is thought to be dependent on many factors, with root growth and sediment anchoring identified as keys to long-term success (31, 68, 69). Despite the importance of these belowground processes, few studies have explicitly examined the impact of microbiome community structure on transplantation success. A study by Milbrandt and colleagues is, perhaps, an instructive exception (28). Similar to our findings, washed and sod transplants of *Thalassia testudinum* showed few differences in plant traits several weeks after transplantation. Critically, though, transplants that were planted into autoclaved sediment demonstrated a strong and significant die-off starting at seven weeks post transplantation, leading the authors to conclude that an intact microbial community is essential to the plant’s ability to combat transplantation shock. An important distinction of our work is that growth traits of washed transplants consistently lagged behind those of sod transplants during the first week of the experiment when microbiome recovery was most pronounced. Given these results and the highly variable nature of restoration outcomes, understanding the roles of the seagrass microbiome in optimizing plant physiology, combating transplantation shock, and contributing to anchoring effects at the bed-scale will be essential to the development of best practices for future seagrass restoration programs.

## Materials and Methods

### Experimental Setup

Sediment (top ∼ 15 cm) and 90 healthy *Z. marina* primary shoots were manually collected at low tide from intertidal eelgrass beds in Yaquina Bay, OR, USA (44.624518, - 124.044372) during July 2018. Sediment was sieved (4 cm^2^) and held in buckets filled with seawater for 24 h. Plants were manually extracted from the beds by excavating a ∼3 cm radius sediment ball around the roots and collecting terminal shoots with attached rhizome fragments, a method that is similar to those previously used in studies on seagrass transplantation (70–72). The loosely attached, non-rhizosphere sediment was dislodged from the rhizome fragment by gentle agitation. This procedure adheres to the operational definition of the rhizosphere -- the sediment attached to the roots after manually shaking (15, 73) -- while also capturing the biological definition of the rhizobiome, i.e., the microbial community that is closely associated with plant roots and is influenced by plant metabolism (1). Plants were placed in plastic bags and processed for transplantation within three hours of collection.

Individual plants were randomly assigned to either the “wash” or “sod” transplant treatment group. The rhizospheres of plants in the washed group were removed by a gentle seawater rinse, replicating the potential rhizosphere loss in transplantation efforts. The rhizospheres of plants assigned to the sod treatment group were left undisturbed. The rhizomes of plants in the wash treatment were trimmed to retain five internodes connected to the first five root bundles (74), and rhizomes of sod transplants were standardized by trimming to lengths matching those of washed plants. Plant leaves were standardized across treatments by trimming to 50 cm (75).

PVC cylinders (18 × 7.6 cm) were filled with sediment and the meristem of each plant was positioned near the top of each. Sediment was added to cover the rhizome, roots, and rhizosphere (if attached). Planters were randomly and evenly placed inside a 2000 liter outdoor flow-through tank filled with water from Yaquina Bay.

### Plant Sampling and Morphometric Analyses

Whole plant sampling was performed on the initial day of the experiment (t = 0) prior to transplantation and on days 1, 3, 7, 14, 21, and 28 post-transplantation. At least five plants from each treatment were collected and destructively sampled at each time point. Plants were initially agitated to remove loosely attached sediment. The rhizosphere sediment was then washed from plant roots in 25 ml of sterile seawater and collected in sterile tubes. One ml of the resulting slurry was transferred to a sterile microcentrifuge tube and stored at -80 °C until DNA extraction. One pair of the youngest root cluster was then removed from the plant, transferred to a sterile microcentrifuge tube, and stored at -80 °C for DNA extraction.

Roots not used for extractions were removed from plants, counted, and measured to calculate average lengths. Rhizome lengths and longest leaf lengths were recorded for plants. Biomass measurements were recorded for the component parts of plants (i.e., leaves, rhizomes, and roots) after drying for seven days at 40 °C. The residual sediment slurries from plants (∼24 ml/plant) were vacuum-filtered through pre-weighed GFF membranes, dried as above, and net weights were recorded as rhizosphere masses.

### DNA extraction, PCR, and Amplicon Sequencing

Microbial community DNA was extracted from frozen roots and sediment slurries using a CTAB and phenol:chloroform extraction method (76) within six weeks of sample collection. Amplicon sequencing libraries were constructed from 25-100 ng of template DNA using a one-step PCR with bar-coded 515F and 806R universal 16S rRNA (v3-v4) primers (77). PCRs were performed using AccuStart II ToughMix Polymerase following the manufacturer’s instructions and performing a thermal cycle program of: 94 °C (3 min.); 25 cycles of 94 °C (45 sec.), 50 °C (60 sec.), 72 °C (90 sec.); 72 °C (10 min.); 4 °C (hold).

Successful amplification reactions (139 of 143 samples) were purified using Agencourt AMPure XP beads following the manufacturer’s instructions, with the exception that a 1:1 ratio of bead solution and PCR product was used. A Qubit 2.0 fluorometer (Thermo Fisher Scientific, Waltham, MA, USA) was used to quantify concentrations of purified amplicons, and these values were used to evenly pool libraries prior to sequencing with the Illumina MiSeq (Illumina Inc., San Diego, CA, USA).

The ‘DADA2’ package (v 1.10.1) (78) within the Bioconductor software environment (v 3.8) (79) of the R Project (v 3.5.2) (80) was used to process raw sequencing reads. All reads were initially quality filtered using the ‘filterAndTrim’ command with default settings (“maxN=0, maxEE=c(2,2), truncQ=2”). To avoid computational limitations resulting from the fact that multiple libraries contained >>100000 reads, the resulting high-quality reads of libraries were randomly down-sampled to 15000 paired-end reads (BioProject ID: PRJNA591021). This resulted in 126 libraries with ≥ 8891 high-quality paired-end reads used as inputs for the remaining DADA2 pipeline (i.e., error-rate training, sample inference, paired-read merging, chimera removal, ASV (Amplicon Sequence Variant) counting, and taxonomic assignment against the SILVA Ref NR 132 database) (81). An average of 7819 ± 1430 sequences were retained across all libraries (Table S1), and sequence counts were rarefied to the library with the minimum count (n = 4881) using the ‘rrarefy’ function of ‘vegan’ (v 2.5-5) (82). A final count table with individual samples containing 119 ± 27 ASVs and 2296 ASVs detected across all samples was generated.

A filtered alignment of representative ASV sequences against the pre-computed SILVA Ref NR 132 alignment was created using the ‘align.seqs’ and ‘filter.seqs’ commands of the mothur software package (v 1.40.5) (83). FastTreeMP (v 2.1.7) (84) calculated a phylogenetic tree from the filtered alignment applying a generalized time-reversible model of evolution (85). The resulting tree was midpoint rooted using ‘reroot.pl’ (86).

### Statistical Analyses

The ‘phyloseq’ package (v 1.26.1) (87) was used to import the phylogenetic tree, count table, taxonomy table, sequence FASTA of ASVs, and a matrix containing plant trait data, sampling date, plant compartment information, and treatment assignments for each sequence library into R. Single pseudocounts were added to plant trait variables containing zeros, allowing for log_2_-transformation. All statistical testing was performed in R and plots were created using ‘ggplot2’ (v 3.1.1) (88) and ‘ggpubr’ (v 0.2.1) (89). Summary statistics are reported as means (M) plus/minus standard deviation, unless otherwise stated.

The ‘vegdist’ function of ‘vegan’ was used to create a Euclidean distance matrix of samples based on log_2_-transformed, centered, and scaled plant morphometric data. The ‘UniFrac’ function of ‘phyloseq’ created weighted UNIFRAC distance matrices (90) from count tables and the phylogenetic tree. To test for the significance of sample clustering, the ‘adonis2’ function of ‘vegan’ was used with 1000 permutations (91). Two- and three-way tests were performed multiple times with the order of the independent variables in the formula changed to ensure consistency of test results, regardless of term precedence. To visualize sample distance relationships, Principal Coordinates Analyses (PCoAs) (92) were performed using the ‘pcoa’ command of ‘ape’ (v 5.3) (93). In figures, percentages on axes labels of PCoA plots report the percent variation captured by each coordinate, and axes lengths are scaled to this number. Spearman’s rank correlations (ρ) between distance matrices of plant trait and ASV count data were determined using the ‘bioenv’ and ‘mantel’ functions of ‘vegan’.

Significant effects of treatment and/or time on response variables were assessed with Student’s T-tests and Analyses of Covariance (ANCOVAs) using the ‘t.test’ and ‘ancova’ functions of ‘stats’ (v 3.5.2) and ‘HH’ (v 3.1-37) (94). If no significant interactions between the treatment effect and the time covariate were detected, an Analysis of Variance (ANOVA) was performed on a reduced model without the interaction term using the ‘Anova’ function of the ‘car’ package and applying Type II sum of squares calculations (95).

Significant differences in ASV abundances between plant compartments (α ≤ .01) were tested using the ‘DESeq’ function of ‘DESeq2’ (v 1.22.2) (96). Generalized linear mixed models (GLMMs) (97) were used to determine significantly different temporal trends in abundance for microbial taxa. A Tweedie compound Poisson distribution was chosen for this model given that it best captures the nature of amplicon sequence datasets (e.g., overdispersion, zero-inflated datasets, and continuous values) (98). The ‘cpglmm’ function of the ‘cplm’ R package (99) was used for time-series analyses following the general procedure outlined in (98). Summarized sequence count tables of family-level taxonomic units were created and full GLMMs were fit relating counts to treatment, days post transplantation, the interaction of main effects, and random effects of each taxon. Taxa detected in > 25% of samples and with cumulative sequence counts > 100 reads were tested to focus on the most abundant, prevalent, statistically robust groups in our samples. *P*-values of modeled slopes and intercepts were obtained via likelihood ratio tests between the full model and two reduced models where the interaction or the treatment variable was removed. Slope and intercept *p*-values were adjusted using the Benjamini-Hochberg method (100), and adjusted values ≤ .05 were considered significant. Resulting intercepts with positive values indicated that a taxon’s initial abundance was higher in washed versus sod transplant rhizospheres, with negative intercepts implying the opposite. Modeled slopes with positive coefficients indicated that rate of increase for a given taxon’s abundance was greater over the course of the experiment in the wash treatment than in sod samples, and vice versa for negative slope coefficients.

## Data Availability

The sequence reads from all samples collected from experiments were deposited in the NCBI data bank (BioProject ID: PRJNA591021). Data access for reviewers only is available at: https://dataview.ncbi.nlm.nih.gov/object/PRJNA591021?reviewer=o44608tibs1lcm4ngr8q8s4f8d

## Acknowledgements

We acknowledge Oregon State University’s Summer Undergraduate Research Experience program, HMSC’s Mamie Markham Research Award, and the Joan Countryman Suit Scholarship for financial support, the CGRB core facility for sequencing support, C. Moffett and I. Cheung at HMSC for facility and administrative support, E. Slick for help with sample collection.

